# Overview of the *Saccharomyces cerevisiae* population structure through the lens of 3,034 genomes

**DOI:** 10.1101/2024.09.16.613241

**Authors:** Victor Loegler, Anne Friedrich, Joseph Schacherer

**Affiliations:** Université de Strasbourg, CNRS, GMGM UMR 7156, Strasbourg, France; Institut Universitaire de France (IUF), Paris, France

## Abstract

With the rise of high-throughput sequencing technologies, a holistic view of genetic variation within populations – through population genomics studies – appears feasible, although it remains an ongoing effort. Genetic variation arises from a diverse range of evolutionary forces, with mutation and recombination being key drivers in shaping genomes. Studying genetic variation within a population represents a crucial first step in understanding the relationship between genotype and phenotype and the evolutionary history of species. In this context, the budding yeast *Saccharomyces cerevisiae* has been at the forefront of population genomic studies. In addition, it has a complex history that involves adaptation to a wide range of wild and human-related ecological niches. Although to date more than three thousand diverse isolates have been sequenced, there is currently a lack of a resource bringing together sequencing data and associated metadata for all sequenced isolates. To perform a comprehensive analysis of the population structure of *S. cerevisiae*, we collected genome sequencing data from 3,034 natural isolates and processed the data uniformly. We determined ploidy levels, identified single nucleotide polymorphisms (SNPs), small insertion-deletions (InDels), copy number variations (CNVs), and aneuploidies across the population, creating a publicly accessible resource for the yeast research community. Interestingly, we showed that this population captures ∼93% of the species diversity. Using neighbor-joining and Bayesian methods, we redefined the populations, revealing clustering patterns primarily based on ecological origin. This work represents a valuable resource for the community and efforts have been made to make it evolvable and integrable to future yeast population studies.

## Introduction

Species naturally exhibit a certain degree of genetic diversity among individuals. This diversity contributes to a significant fraction of observed phenotypic variation and is crucial to both the survival and evolutionary potential of a species. In fact, it has long been established that genetic diversity provides the raw material on which natural selection can act (Fisher 1930). The advent of whole-genome sequencing at the population level has facilitated the analysis of intraspecific genomic diversity, giving rise to the field of population genomics. Short-read sequencing of large populations allows for the construction of dense maps of the genetic polymorphisms, including single nucleotide polymorphisms (SNPs), small insertion-deletions (InDels), and copy number variations (CNVs). Subsequently, analyses of these polymorphisms can provide insights into external forces that shape genomes, such bottlenecks or selection occurring during domestication processes as well as environmental changes.

The budding yeast *Saccharomyces cerevisiae*, given its small genome, high genetic diversity, and complex domestication history, has emerged as a widely used model organism for population genomics. This species has colonized a wide range of wild and human-related habitats and, as a result, has been subject to multiple evolutionary constraints. Isolates have been found in primary and secondary forests in Asia, in fermentation processes of several beverages, including wine, beer, sake and tequila, and have also been identified as opportunistic human pathogens (McCusker et al. 1994; Wang et al. 2012; Parapouli et al. 2020; Gallegos-Casillas et al. 2024). A multitude of population genomics analyses have permitted the description of wild and domesticated populations at the genetic level. Initial studies on a small number of individuals (between 36 and 63) revealed the existence of well-defined populations corresponding to different ecological origins, with the notable identification of wine and sake clades (Liti et al. 2009; Schacherer et al. 2009). The grouping of geographically diverse wine isolates in a single clade, coupled with the low diversity of this clade, points to a single domestication event associated with a bottleneck as the probable origin of all wine isolates. In addition to clades originating from a single ancestry, signals of high admixture were detected in some isolates. Subsequent analysis of 100-genomes with a particular focus on clinical isolates revealed that they are overrepresented in admixed isolates, suggesting that outcrossing facilitated adaptation to this novel ecological niche (Strope et al. 2015). Using the same dataset, regions of high diversity were identified as introgressions from the sister species of *S. cerevisiae*, namely *Saccharomyces paradoxus*. A high number of introgressed genes were reported in isolates that were subsequently assigned to the Alpechin lineage, derived from olive oil production wastewater (Peter et al. 2018; Pontes et al. 2019). To better characterize populations found in human-related niches, a later survey focused on 157 industrial isolates, revealing hallmarks of domestication (Gallone et al. 2016). Notably, beer isolates exhibit high ploidy level, high heterozygosity and impaired sexual reproduction. Further work on Asian isolates demonstrated a single domestication event for all Asian fermentation isolates (Duan et al. 2018). Moreover, wild isolates predominantly found in Asia display a greater level of diversity and a lower heterozygosity compared to domesticated isolates. Efforts were made to assemble and sequence a large collection of 1,011 diverse natural isolates, allowing the precise definition of clades within the species (Peter et al. 2018). This revealed a single out-of-China origin for *S. cerevisiae*, followed by a complex history of domestication and adaptation to a wide range of niches, shaped by introgressions and loss of heterozygosity events. Additional niche-specific populations were later sequenced, with a notable addition of wild isolates sampled in Taiwanese forests (Lee et al. 2022). Although more than 3,000 isolates have now been sequenced, the datasets are scattered and a comprehensive reference of all studies and data is not available. While populations larger than the 1,011 were gathered to investigate the gene-based pangenome of the species (Li et al. 2019; M. Wang et al. 2024; H. Wang et al. 2024) or to focus on introgressions (Tellini et al. 2024), no further refinement of the population structure and the evolutionary history has been achieved.

To conduct a comprehensive analysis of the population structure of *S. cerevisiae*, we collected genome sequences for 3,034 natural isolates and processed the data in a standardized manner. We inferred ploidy, detected SNPs, InDels, CNVs and aneuploidies in the population to build a publicly available resource for the yeast community. Neighbor-joining and Bayesian approaches were applied to redefine the clades, which showed clustering primarily based on ecological origin. By providing the raw gvcf files and standard protocols, we aim to make this resource easily expandable as new isolates are sequenced.

## Materials and methods

### Data collection

We gathered Illumina sequencing data of 3,039 *Saccharomyces cerevisiae* natural isolates, coming from 29 publications, with a minimum sequencing depth of 20X. Reads were mapped on the reference genome using bwa-mem2 v2.2.1 (M. Vasimuddin et al. 2019) with default parameters and samtools sort v1.15.1 (Danecek et al. 2021).

### SNPs and InDels calling

A multi-sample VCF containing both variant and non-variant positions was generated following the GATK Germline short variant discovery workflow (https://gatk.broadinstitute.org/hc/en-us/articles/360035535932-Germline-short-variant-discovery-SNPs-Indels-), with version 4.2.3.0 (Poplin et al. 2018 Jul 24). The raw VCF was filtered with bcftools v1.15.1 (Danecek et al. 2021) for quality (DP >= 10, GQ >= 20), missing genotypes (missing genotype per sample below 20% and missing genotype per loci below 1%) and excess of heterozygosity (ExcHet > 0.99). Out of the 3,039 samples, 3,034 passed the missing genotype filters and were considered for the further analyses. Variants calls were further separated in two files, containing SNPs and InDels, along with non-variant positions. Complex loci, spanning both SNPs and InDels were filtered out. All further VCF filtering was performed with bcftools, unless otherwise mentioned. Callable sites mentioned in the results section refer to the non-variant positions in addition to SNP loci (loci where InDels or both InDels and SNPs are present were not considered for the SNP analysis).

### Zygosity and ploidy estimation

For each sample, the number of heterozygous SNPs was estimated by first extracting each sample from the multi-sample SNPs VCF (bcftools view -s), then filtering only heterozygous positions (bcftools view -i ‘GT=\”het\”‘). Heterozygosity was computed as the ratio of heterozygous SNPs over the total number of callable sites in the SNPs VCF (9,368,983 positions). Samples containing more than 500 heterozygous SNPs (1,756 samples) were considered as heterozygous, the rest were considered homozygous (1,278 samples).

For samples coming from Peter *et al*. 2018, Lee *et al*. 2022 and Marr *et al*. 2023 (1,207 samples in total), ploidy information was retrieved from the original papers as it was estimated experimentally using flow cytometry. For the remaining samples, ploidy was estimated from sequencing data using nQuire (Weiß et al. 2018). Samples with known ploidy were used to benchmark each way of estimating the ploidy with nQuire: (i) maximizing the log likelihood with nQuire lrdmodel, (ii) using the lower SSR or (iii) higher regression R2 of the nQuire histotest command. The latter had the best true positive rate and was used to estimate ploidy of the remaining samples. As ploidy cannot be reliably estimated for homozygous samples, those sample’s ploidy is set to unknown (648 samples).

### CNV calling and aneuploidy detection

CNVs along the genome were estimated for each sample with a depth-based method, CNVnator v0.4.1 (Abyzov et al. 2011), using a 1kb window size. CNV of each CDS was computed as the median normalized sequencing depth of the CDS. In practice, for each position of the reference genome, we assigned a normalized sequencing depth of either 1 when the region was not detected as a CNV, or the value of the Normalized_RD column of the CNVnator output. The median normalized sequencing depth was then calculated for each CDS. A chromosome was considered as aneuploid when more than half of its length was in CNV.

### Clades definition

Clades were defined with a hybrid method using fastStructure (Raj et al. 2014) and manual refining based on the neighbor-joining tree. fastStructure was run on SNPs filtered for biallelic variants, minor allele frequency superior to 5% and LD-pruned with plink 1.9 (Chang et al. 2015) (plink --indep-pairwise 50 1 0.5), corresponding to a total of 25,194 SNPs. fastStructure was run with K from 2 to 50. The chooseK command was used to identify the best number of components in the population (Model components used to explain structure in data: K = 38). The neighbor-joining tree was constructed with 1,918,693 SNPs using the R packages SNPRelate (Zheng et al. 2017) and ape (Paradis and Schliep 2019). 38 groups were constructed based on the K=38 ancestry components, taking samples with more than 60% of ancestry. Groups were further manually refined to remove outliers on the neighbor joining tree. Groups with unclear segregation on the tree (within wine isolates) were merged and manually clustered based on the phylogeny. Despite being in the same ancestry component, Mexican agave and French Guiana samples were split in 2 groups because of their ecological origin difference and their clear separation on the tree. 39 clades were finally obtained. Four superclades were then constructed by grouping clades according to concordant ecological origin and phylogeny.

## Results

### Species-wide diversity

To create a comprehensive set of sequenced isolates, we filtered public sequence databases for Illumina whole-genome sequences of *S. cerevisiae* strains, harboring a minimum sequencing depth of 20X. All genetically modified isolates were discarded. In the end, we collected whole-genome sequencing data for a total of 3,039 isolates of *S. cerevisiae* generated through 29 population genomics surveys (Dunn et al. 2012; Zheng et al. 2012; Almeida et al. 2015; Hose et al. 2015; Marsit et al. 2015; Song et al. 2015; Strope et al. 2015; Barbosa et al. 2016; Borneman et al. 2016; Franco-Duarte et al. 2016; Gallone et al. 2016; Gayevskiy et al. 2016; Zhu et al. 2016; Coi et al. 2017; Kita et al. 2017; Maclean et al. 2017; Duan et al. 2018; Peter et al. 2018; Preiss et al. 2018; Fay et al. 2019; Basile et al. 2021; Han et al. 2021; Higgins et al. 2021; Ruiz et al. 2021; Lee et al. 2022; Marr et al. 2023; Morard et al. 2023; Ward et al. 2024). The isolates were sampled in 94 countries across six continents (Fig. 1A) and have a large variety of ecological origins. Isolates are involved in many human-related processes, including winemaking, beer fermentation, spirits production, bakery, in addition to clinical and wild isolates (Table S1). The sequencing reads of all 3,039 isolates were mapped on the reference genome to infer the genetic variants in the population. A total of 1,918,693 SNPs and 58,947 InDels were detected across 3,034 isolates, and five were discarded because of a high fraction of missing genotypes. SNPs account for 20.5% of callable sites on the reference genome. The population nucleotide diversity (median π = 3.6×10^−3^) is slightly increased from the previous estimations based on 1,011 isolates (Peter et al. 2018). We find a mean number of 51,924 ± 22,907 SNPs (0.6% of callable sites) between two randomly selected isolates (Fig. 1B), and a maximum number of 168,614 SNPs (1.8%) between a wild Taiwanese isolate (CEI) and a Mexican isolate from agave distillation (XTRA_FIU). The level of heterozygosity within the population is highly variable, with 1,278 (42%) isolates being homozygous and others having up to 65,785 heterozygous SNPs. Ploidy information based on flow-cytometry data was retrieved from the original publication for 1,207 isolates. For the rest of the population, the ploidy of heterozygous isolates was estimated from allelic frequencies. The ploidy of homozygous isolates without flow cytometry data (648 isolates) could not be retrieved solely from sequencing data, leading to a total of 2,386 isolates with ploidy information. Although most of the isolates are diploid (74% of isolates with known ploidy), a large variation in ploidy level is observed within the population. Polyploids, having more than two copies of the genome, account for 20% of the isolates, and 6% are haploid. Expectedly, the rate of heterozygosity increases with the number of genome copies (Fig. 1C, ρ = 0.5767823, *P* value < 2.2×10^−16^). Seven haploid isolates present a considerable number of heterozygous sites (from 666 to more than 4,500 heterozygous SNPs), because of the presence of aneuploidies or large segmental duplications in their genomes. Aneuploidies were detected based on chromosomal sequencing depth for each isolate and are frequent in the population, with 927 isolates (31%) having at least one aneuploid chromosome. As for the heterozygosity rate, we observe an increase of the frequency of aneuploidies in isolates with a high ploidy (Fig. 1D). While 28% of the diploid isolates are aneuploid, this proportion rises to 52% for polyploid isolates. The frequency of aneuploidies is also correlated to the length of the chromosome, as 74% of the observed events affect the 4 shortest chromosomes (chromosome 1, 3, 6 and 9) (Fig. S1).

**Fig. 1.**
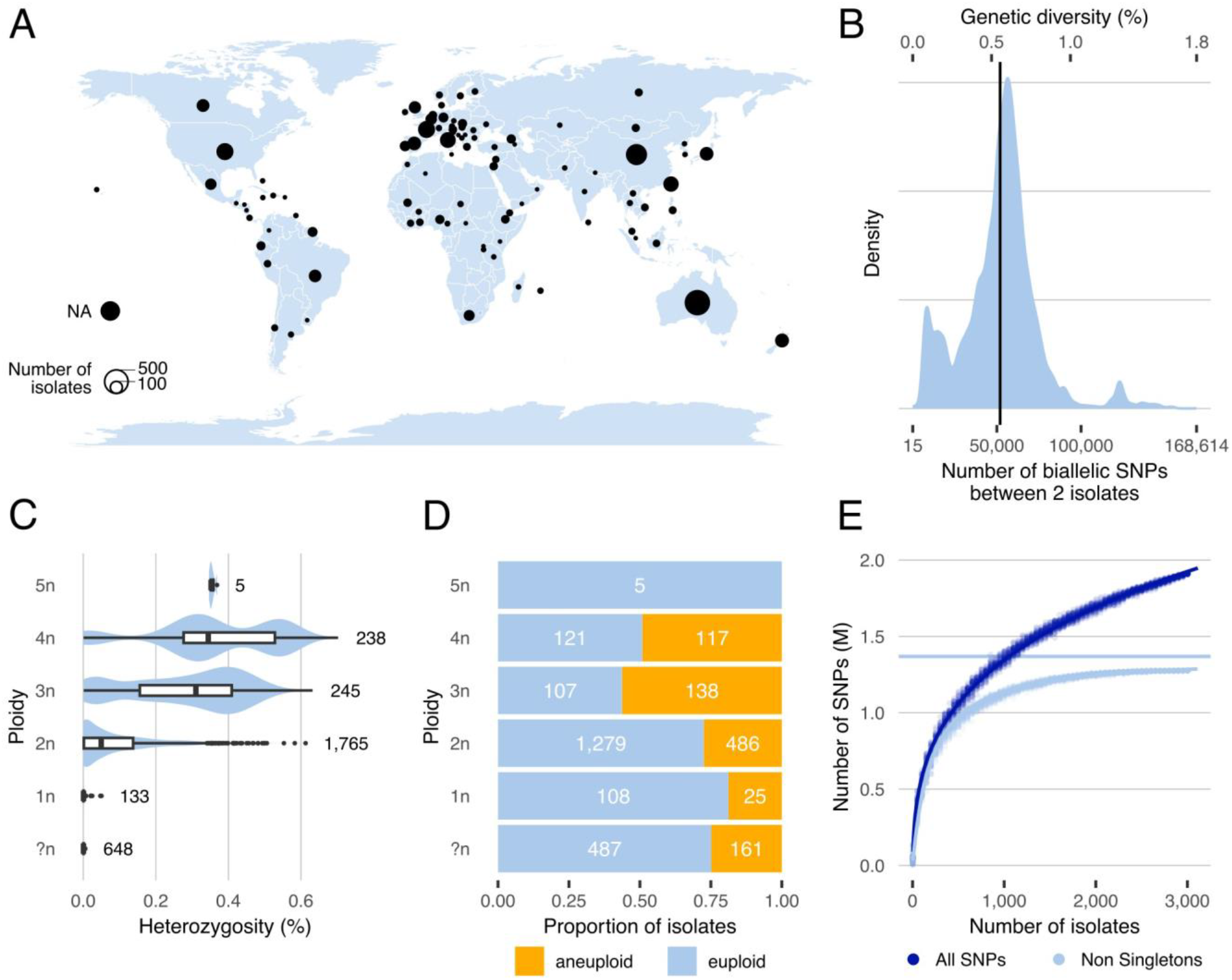
Origin and diversity of 3,034 natural isolates. **A**. Map of the geographical origin of the isolates. **B**. Distribution of the number of SNPs between two randomly selected isolates. The vertical line represents the mean number of 51,925 SNPs. The percentage of genetic diversity is the ratio of the number of sites to the number of callable sites. **C**. Percentage of heterozygosity for each ploidy level. Heterozygosity is the ratio of the number of heterozygous SNPs to the number of callable sites. **D**. Number of euploid and aneuploid isolates per ploidy level. **E**. SNPs rarefaction curves, using all SNPs (dark blue) and non-singleton SNPs (light blue). The curve for all SNPs was fitted with a power law curve (*y* = 242,965 × *x*^0.276^ − 288,549), and the curve with non-singleton SNPs with a Michaelis-Menten equation (*y* = (1,369,438 × *x*)/(204 + *x*).

### Estimation of the species diversity captured by the 3,034 population

To estimate the fraction of species diversity captured by the population, we built rarefaction curves of the genetic variations in the population using SNPs and InDels as markers, taking all variants and non-singleton variants (*i*.*e*., variants present in at least two isolates). The increase in the number of variants as a function of the number of isolates follows a power-law, making extrapolation impossible as the number of variants tends towards infinity (Fig. 1E, Fig. S2). However, since the number of non-singleton variants follow a Michaelis-Menten equation, it is possible to estimate the total number of non-singleton variants in the species as the limit of the equation with an infinite number of isolates. Accordingly, our population covers 93.2% of non-singleton SNPs and 92.9% of non-singleton InDels in the species. In comparison, the 1,011 *S. cerevisiae* collection (Peter et al. 2018) covers 60.8% of the species diversity (SNPs-based, 60.9% using InDels). It should be noted that this estimation corresponds to the higher bound of species diversity fraction covered by our dataset. The addition of a new divergent population would considerably increase the number of non-singleton variants, thereby reducing the diversity captured by the 3,034 isolates.

### A broader view of *S. cerevisiae* population structure

We sought to redefine the structure of the *S. cerevisiae* population by constructing a neighbor-joining tree of the 3,034 isolates based on 1,918,693 SNPs. In addition, population structure was inferred based on 25,194 common and pruned SNPs, using an ideal number of K=38 ancestry components to explain the population structure (Fig. S3). Both approaches were combined and manually refined to define 39 clades within the population (Fig. 2, Fig. S4, Tables S1-2). The clades were named based on the geographical or ecological origin of the majority of the constituent isolates. Additionally, four superclades were defined by grouping clades with similar ecological origin and consistent position on the phylogenetic tree. Thus, the Wine (1,247 isolates), Beer (310), Asian Fermentation (305) and Wild (343) superclades were defined (Fig. 2). The number of isolates per clade ranges from nine (32. US clinical 3) to 273 (8. AU Wine 4), with a median of 46 isolates per clade. A total of 321 isolates (10.6%) were not assigned to any clade or superclade due to their admixture of multiple ancestry components. We observe a sampling bias toward domesticated isolates, with 2,177 isolates, compared to wild (390 isolates) and other (146 isolates) ecological origins. In comparison to the 1,011 *S. cerevisiae* dataset (Peter et al. 2018), the 3,034 population contains novel genetic clades as well as a globally increased sample size. The collection mainly gained Australian wine isolates (Ward et al. 2024), which form specific clades diverging from the previous wine clades, and Asian fermentation isolates (Duan et al. 2018; Han et al. 2021) that add novel diversity around the sake clade. These additional clades explain the increase in nucleotide diversity of the global population. There is also a substantial increase in the sample size of Chinese and Taiwanese wild isolates, although the diversity brought by these clades were already represented in the 1,011 population (Lee et al. 2022). A notable increase in the sample size of Wine superclade now permits the observation of the Alpechin clade branching within the wine isolates (Fig. 2, Fig. S4). This suggests that the Alpechin lineage originated from domesticated wine isolates that colonized this niche and hybridized with *S. paradoxus* isolates.

**Fig. 2.**
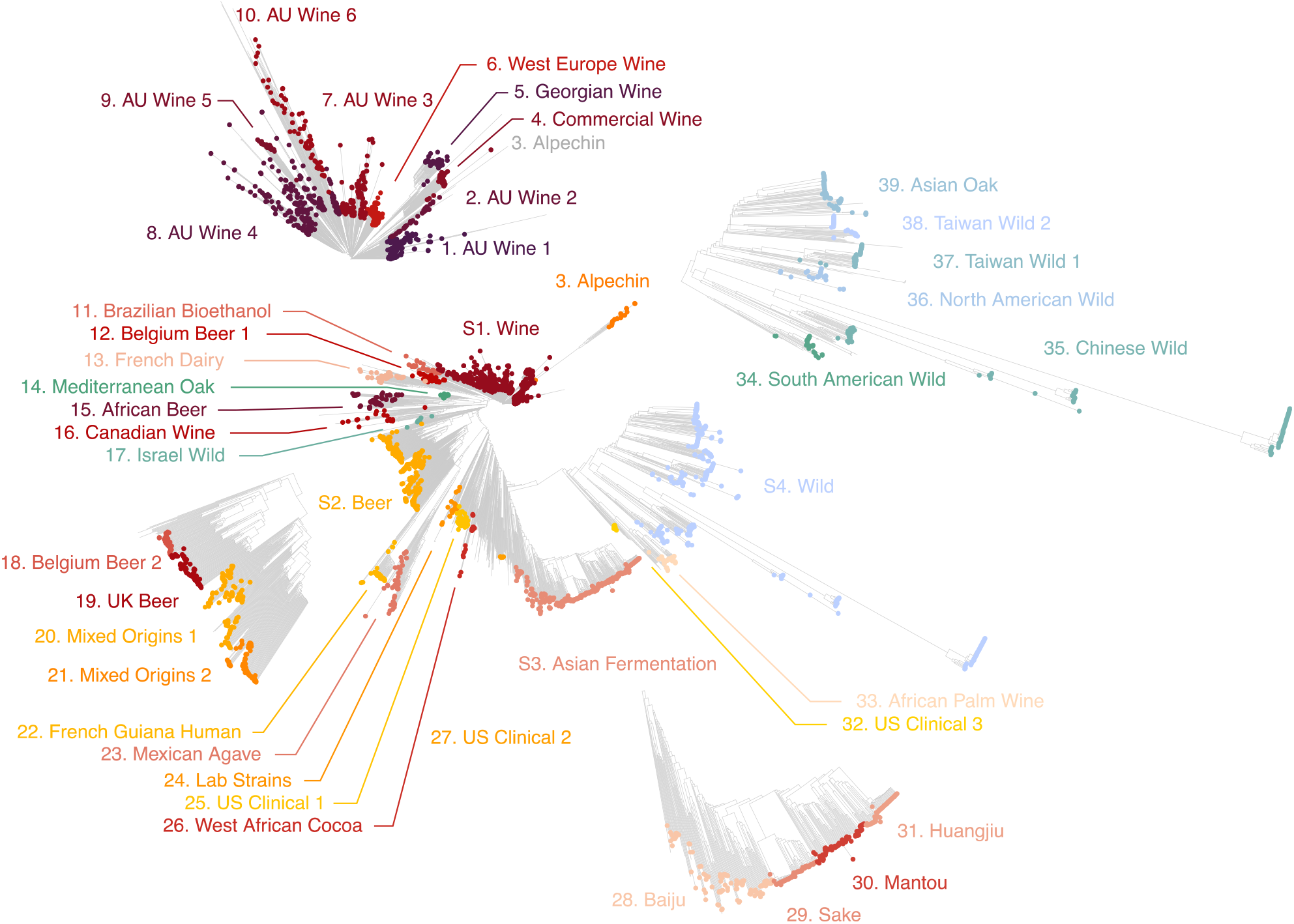
Neighbor-joining tree built using 1,918,693 SNPs. Isolates are colored according to the 39 clades and 4 super-clades that were identified. Subtrees of the four superclades are magnified (S1. Wine, S2. Beer, S3. Asian Fermentation and S4. Wild).

### Differences in diversity between clades

A variable level of diversity is observed across clades. Notably, the diversity observed at the ploidy level is highly clade dependent. Polyploid isolates are almost exclusively found in the Beer and Asian Fermentation superclades, suggesting that polyploidization may be a hallmark of these domestication processes (Fig. 3A). An enrichment for polyploid isolates is significant in Beer isolates (odds ratio = 0.13, *P* value < 2.2×10^−16^), as previously described, but also in Baiju isolates, where 35% are polyploids (odds ratio = 0.34, *P* value = 8.7×10^−7^). However, despite domestication, the Wine superclade exhibit a similar proportion of polyploids compared to Wild isolates (odds ratio = 1.6, *P* value = 0.15). In addition to the reported variation in ploidy, a variation in heterozygosity level is also observed. The heterozygosity of Beer isolates is significantly higher than that of the rest of the population (W = 794969, *P* value < 2.2×10^−16^), with this difference driven by a positive correlation between high ploidy and high heterozygosity rate (Fig. 3B, Fig. 1C). To a lesser extent, a similar increase in heterozygosity is observed in Asian Fermentation isolates, driven by Baiju polyploid isolates (W = 245440, *P* value < 2.2×10^−16^). Nevertheless, the prevalence of polyploid isolates does not fully account for the elevated heterozygosity observed in domesticated clades. Despite a comparable proportion of polyploid isolates, the heterozygosity rate in the Wine superclade is slightly higher than that of Wild isolates (W = 224532, *P* value = 0.022). This indicates that heterozygosity may be a second hallmark of domestication. Finally, a variation in the pairwise nucleotide diversity is also evident across clades. Despite the increased sample size for Wine isolates, the Wine superclade still exhibits the lowest diversity, providing evidence for the strong bottleneck that occurred during the domestication of Wine isolates (Fig. 3C). While Wild isolates have a greater diversity globally than domesticated clades, isolates of the Chinese Wild clade exhibit particularly high diversity compared to the other clades. This clade, previously defined as the Taiwanese clade, has been reported as the ancestral wild *S. cerevisiae* population (Peter et al. 2018).

**Fig. 3.**
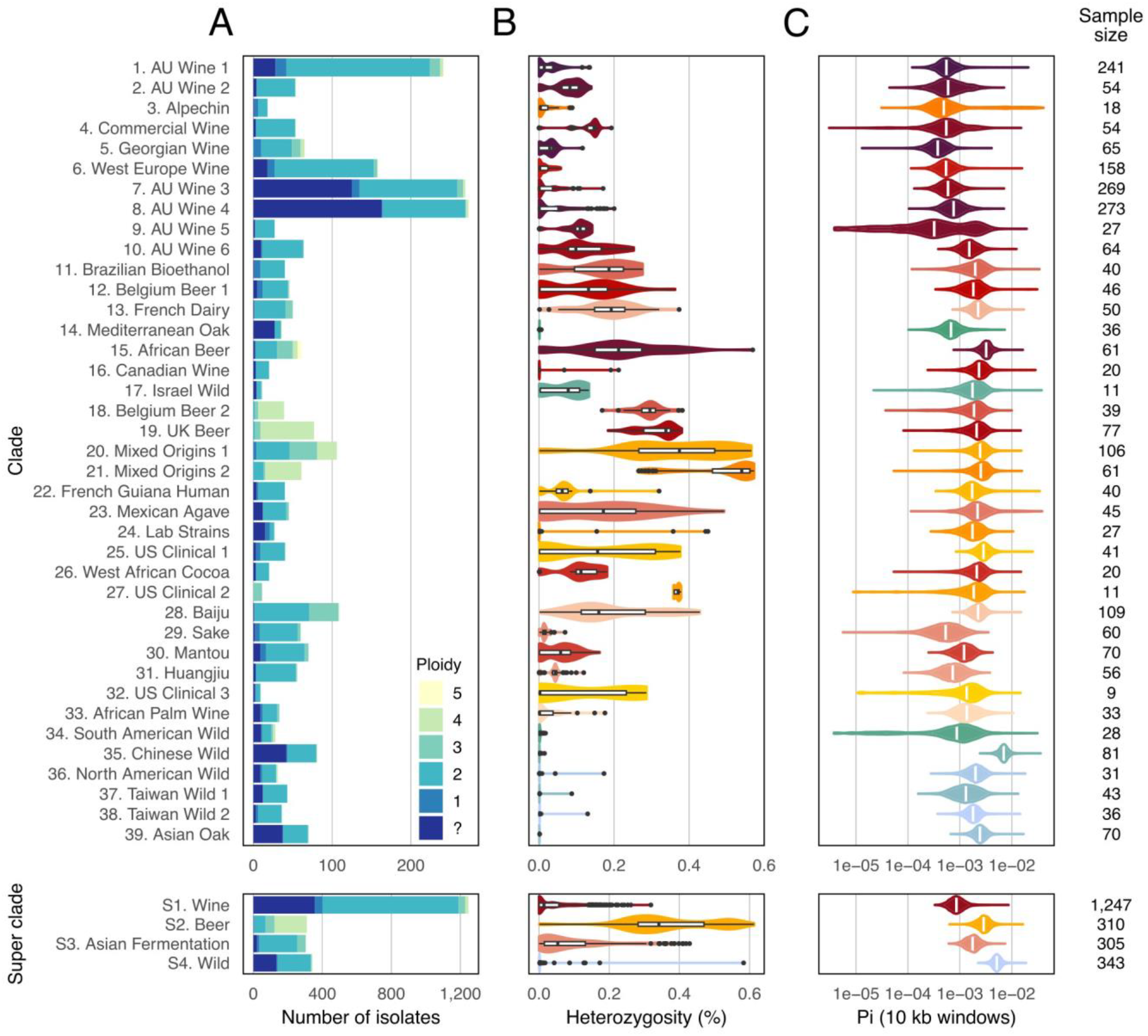
Diversity per clade. **A.** Ploidy level per clade. **B**. Fraction of heterozygous position per clade. **C**. Pairwise nucleotide diversity per clade, computed over 10 kb windows sliding by 1 kb steps.

### Creation of a shareable resource

In addition to providing an overview of the natural population of *S. cerevisiae* sequenced so far, this work aims to build an accessible resource that catalogs sequenced isolates as well as natural genetic variants in the species. The metadata for the whole population was gathered, including geographic and ecological origin (Table S1). Additionally, zygosity, ploidy (when available) and aneuploidies were systematically referenced for each isolate (Table S1). To catalog small genetic variants, we used a standardized and commonly used variant calling pipeline (see Methods) and provide catalogs of high-quality SNPs and InDels (Loegler et al. 2024a). Furthermore, CNV data were estimated for each gene of the reference genome in each isolate. As there is no standard way to detect and to encode CNVs, we decided not to provide a number of gene copies, but rather give a normalized sequencing depth for each gene (see Methods). By providing minimally processed data, we hope to prevent any potential bias that may arise from choosing an arbitrary depth threshold to state the presence of a gene. Future users will be accountable for transforming this data according to their own preferences. This catalog of small variants and CNVs will provide information about occurrence and frequency of natural variants to the yeast community.

Moreover, the inclusion of all or part of 3,034 isolates in future genomic studies may be highly valuable to locate newly sequenced isolates within the species phylogeny. However, processed variant matrices cannot be properly combined with novel sequencing data, as the sequencing data for each isolate must be handled in the exact same way. Collecting and processing raw sequencing reads from a previously sequenced large population can be extremely time-consuming, as sequencing files are large (more than 5 terabytes for the 3,034 isolates) and scattered across databases. To facilitate the proper integration of the 3,034 population in future genomic analyses, we provide the gvcf file of each isolate, which can be combined with novel sequencing data to build a multisample vcf (Loegler et al. 2024b). A detailed protocol will make this operation straightforward to implement (https://haploteam.github.io/ProtocolSacePopulation).

## Conclusion

Using the sequenced genomes publicly available for more than 3,000 isolates of *S. cerevisiae*, we established a large catalog of the small variants and CNVs of the species and redefined the clades present in this population. Our dataset exceeds by more than three times the number of isolates used for the previous structure analysis (Peter et al. 2018). We notably highlight three domesticated superclades, that include Wine, Beer, and Asian fermentation isolates, and one wild superclade. While the wild superclade is mainly originating from Asia, as previously known, the presence of isolates from North and South America within the same genetic cluster suggest a broader geographic distribution of this clade. Further effort in sampling wild isolates across the world is required to confirm this observation. The differences in ploidy, heterozygosity and nucleotide diversity of the domesticated superclades suggest unique domestication processes, and considering all domesticated isolate as a single group for comparison with wild isolates is not appropriate. While many studies argue on the possibility of multiple domestication events for *S. cerevisiae* (Fay and Benavides 2005; Schacherer et al. 2009; Gallone et al. 2016; Peter et al. 2018), further research is necessary to gain a precise understanding of the demographic history of the species. To address this question, our dataset provides the right material to perform complex demographic modeling (Schraiber and Akey 2015).

Considering the multiplicity of *S. cerevisiae* sequencing datasets and the difficulties in collecting them, we aim to establish this work as a valuable resource for the yeast community by giving access to the catalog of genetic variants and metadata for this population. As the number of sequenced isolates keeps increasing, we also provide the raw gvcf files that can be used to build the multisample variant matrices, along with a standardized protocol to process the reads (https://haploteam.github.io/ProtocolSacePopulation). This will facilitate the inclusion of all or part of the 3,034 population in future population genomic studies.

A major limitation of our study stands in the type of genetic variation detected. The use of short-read sequencing data does not allow for the reliable detection of structural variants (SVs) in the population, which is essential for the generation of a comprehensive catalog of genetic variants at the population level. Moreover, the variation in the gene content can only be partially assessed, as contiguous genome assemblies are needed for the proper construction of a gene-based pangenome. In light of this, long-read sequencing has been conducted in *S. cerevisiae* on smaller populations, offering a preliminary description of the SV landscape (Istace et al. 2017; Lee et al. 2022; O’Donnell et al. 2023). However, given the limited sample size, the SVs detected did not reach saturation, leaving a fraction of the genetic diversity uncaptured. By extrapolating the results obtained from telomere-to-telomere genome assemblies of 142 isolates (O’Donnell et al. 2023), it can be assumed that a population of at least 500 individuals would be needed to catalog SVs in an exhaustive manner. The combination of high-quality genome assemblies for such a population and the dataset we have assembled will provide a comprehensive characterization of the genetic variation present within the species.

## Data availability

Codes used for all analyses and figures, and catalogs of SNPs, InDels and CNVs are available in a first zenodo repository: https://doi.org/10.5281/zenodo.12580561 (Loegler et al. 2024a). GVCF files are available in a second repository: https://doi.org/10.5281/zenodo.12571280 (Loegler et al. 2024b).

First zenodo repository: VCF files

- full3039Matrix.AllPositions.DP10.GQ20.Mind20.99pNonMiss.ExcHet99.INDELs.Var.vcf.gz
- full3039Matrix.AllPositions.DP10.GQ20.Mind20.99pNonMiss.ExcHet99.SNPs.Var.vcf.gz
- Sace_S288c_reference_FullMatrixID.fna
- Sace_SGD_R64-4-1_20230830.gff
- full3039Matrix.GeneNormSeqDepth.tsv.gz
- full3039Matrix.CNVnatorResults.tsv.gz
- Scripts.tar.gz
- Figures.tar.gz

Temporary link:

https://zenodo.org/uploads/12580561?token=eyJhbGciOiJIUzUxMiJ9.eyJpZCI6IjUxZTczNTQ1LWU1ZDEtNDU0Yy05NDQ5LTZjMTc4NmI5OTA1YSIsImRhdGEiOnt9LCJyYW5kb20iOiJiM2ZmYjVmZmFiZmU0OGQ4NjhlZDZiMDFkZWMxNTljYSJ9.RO0XMUdW7nGMGuAvpfecGutvTKRJkDY-ROH3WMwKmYyKcEDxQ2sSN_mgGI3

Second zenodo repository: GVCF files (45G)

- GVCF_3039samples.tar.gz
- Sace_S288c_reference_FullMatrixID.fna
- Sace_S288c_reference_FullMatrixID.bed
- Protocol.pdf

Temporary link:

https://zenodo.org/uploads/12571280?token=eyJhbGciOiJIUzUxMiJ9.eyJpZCI6IjJmMDhhYmFiLWRiOGEtNDkzNy04MzZkLWRiNTM0NWVkMTkwYyIsImRhdGEiOnt9LCJyYW5kb20iOiI2NWYzMGZkZTQyMDc4MmMwMzI0M2I5MTIzYmVlOTA2NSJ9.-v1lFXRylerfgofl0vciLu-hLCV1YwOfCYQvNvlqpB-55vwiPnbsHF1qi7X

## Supplementary figures

**Fig. S1.**
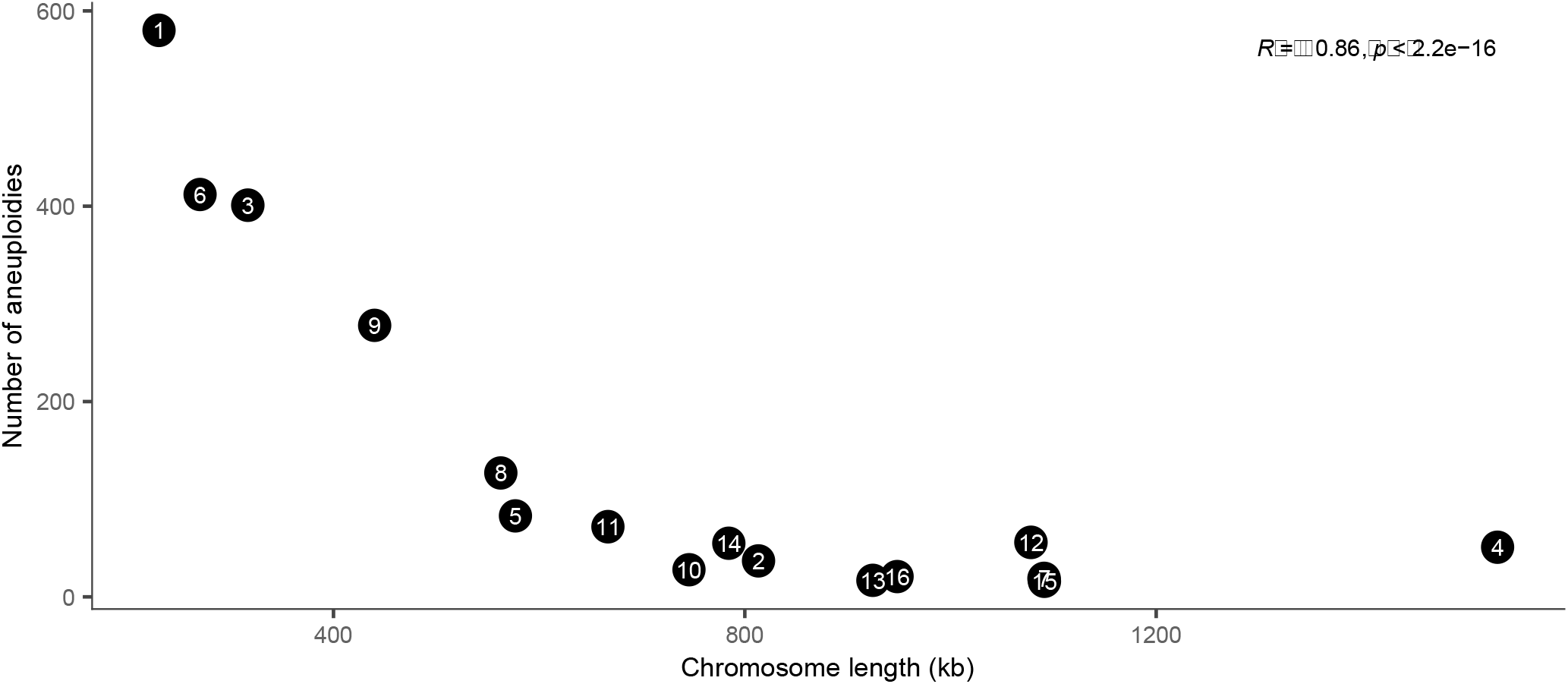
Correlation between the chromosome length and the number of aneuploidies found in the 3,034 population.

**Fig. S2.**
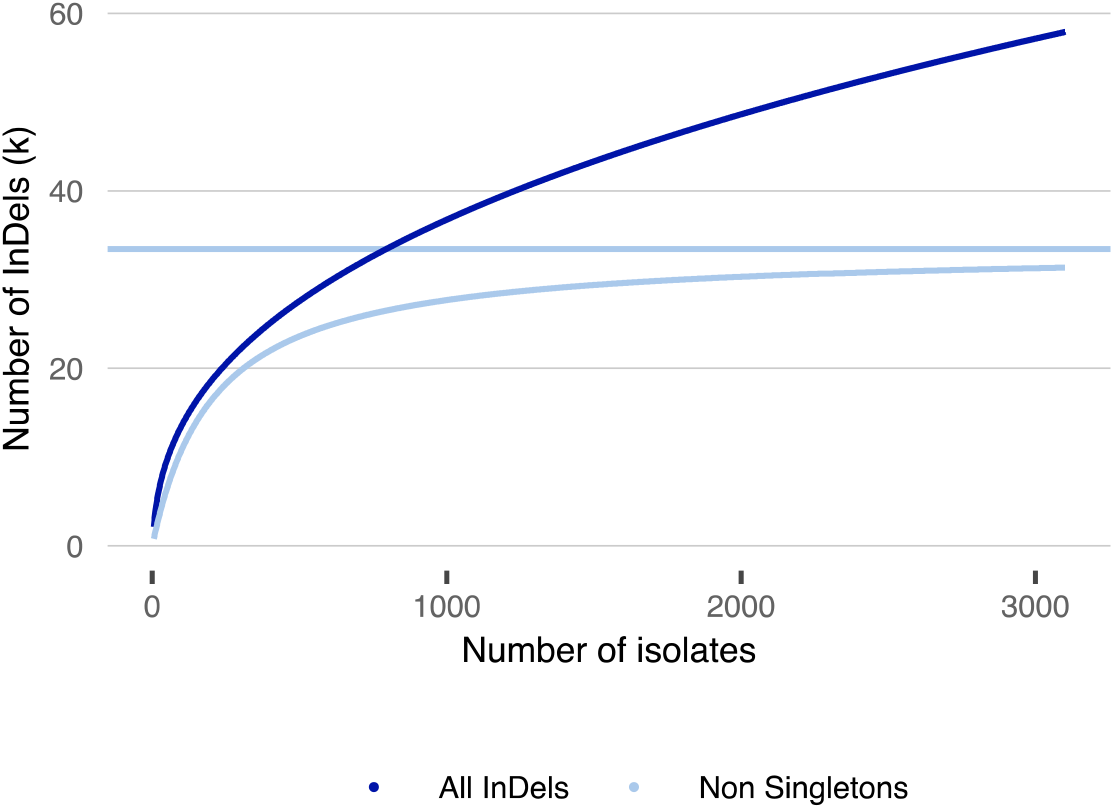
InDels rarefaction curves. Both all InDels (dark blue) and non-singleton InDels (light blue) were used. The curve for all InDels was fitted with a power law curve (*y* = 3,027 × *x*^0.374^ − 3,377), and the curve with non-singleton SNPs with a Michaelis-Menten equation (*y* = (33,450 × *x*)/(208 + *x*)

**Fig. S3.**
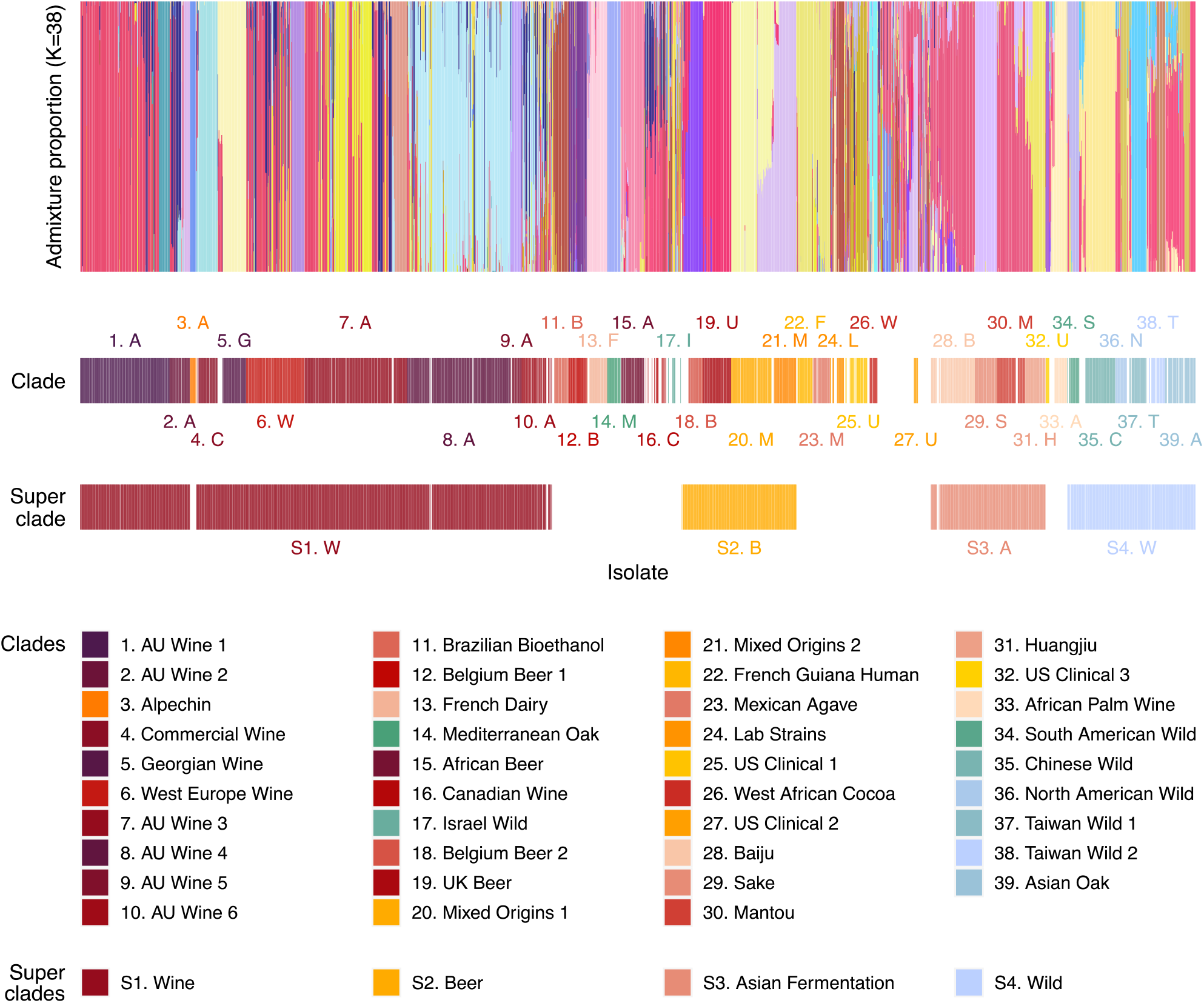
Structure plot of the 3,034 population. The structure plot is represented for the chosen value of K=38. The clade and superclade attributed to each isolate are indicated below.

**Fig. S4.**
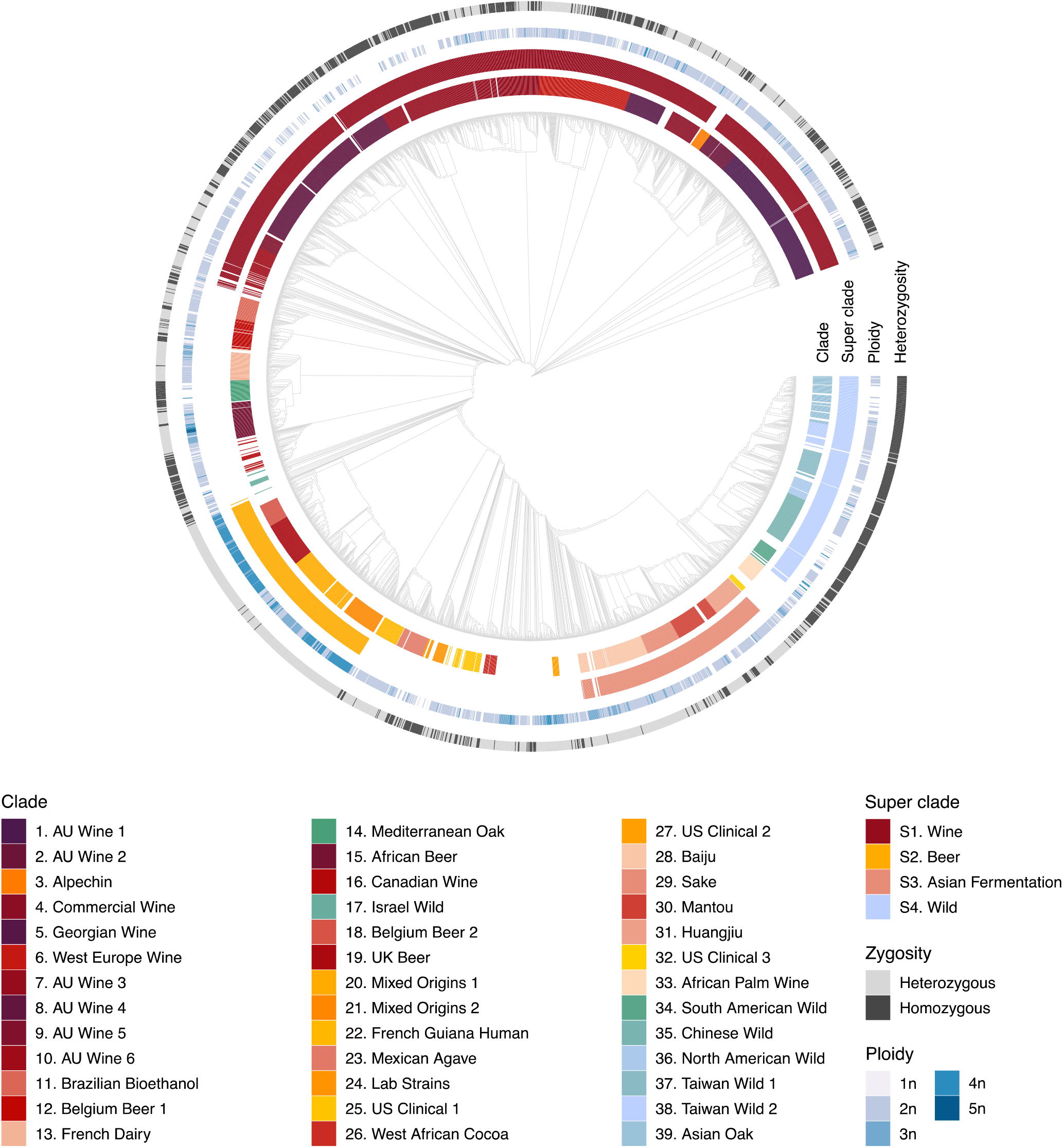
Neighbor-joining tree with clade, zygosity and ploidy information. The neighbor-joining tree was built using biallelic SNPs and is represented without distance information. Isolates are colored according to the 39 clades and 4 superclades that were identified. Zygosity and ploidy information are indicated for each strain. Heterozygosity is defined here as more than 500 heterozygous SNPs per isolate.

